# *dms-viz*: Structure-informed visualizations for deep mutational scanning and other mutation-based datasets

**DOI:** 10.1101/2023.10.29.564578

**Authors:** William W. Hannon, Jesse D. Bloom

## Abstract

Understanding how mutations impact a protein’s functions is valuable for many types of biological questions. High-throughput techniques such as deep-mutational scanning (DMS) have greatly expanded the number of mutation-function datasets. For instance, DMS has been used to determine how mutations to viral proteins affect antibody escape (Dadonaite et al. 2023), receptor affinity (Starr et al. 2020), and essential functions such as viral genome transcription and replication (Li et al. 2023). With the growth of sequence databases, in some cases the effects of mutations can also be inferred from phylogenies of natural sequences (Bloom and Neher 2023) (Figure 1).

The mutation-based data generated by these approaches is often best understood in the context of a protein’s 3D structure; for instance, to assess questions like how mutations that affect antibody escape relate to the physical antibody binding epitope on the protein. However, current approaches for visualizing mutation data in the context of a protein’s structure are often cumbersome and require multiple steps and softwares. To streamline the visualization of mutation-associated data in the context of a protein structure, we developed a web-based tool, `dms-viz`. With `dms-viz`, users can straightforwardly visualize mutation-based data such as those from DMS experiments in the context of a 3D protein model in an interactive format. See https://dms-viz.github.io/ to use `dms-viz`.

## Statement of Need

We wanted `dms-viz` to provide the following functionalities:

1. **Provide structural context**: The main objective of `dms-viz` is to simplify the process of visualizing mutation data with structural context by superimposing mutation measurements on a 3D protein structure. Additionally, it provides extensive control over the visual representation of the 3D structure.
2. **Accommodate diverse data types**: Although analyzing DMS data is a key goal of `dms-viz`, there are many types of mutation data. The tool can handle diverse data types via a command line interface that simplifies the process of converting data into a common format for analysis.
3. **Display multiple conditions**: With `dms-viz`, multiple experimental conditions can be visualized concurrently, facilitating comparisons. Researchers can, for instance, easily visualize deconvolved antibody binding footprints from polyclonal sera (Yu et al. 2022).
4. **Maximize customizability**: Every dataset has specific needs for visual representation. Recognizing this, `dms-viz` offers a high level of customizability. Users can tailor filters, which are important for navigating large and possibly noisy datasets, and tooltips, ensuring that the nuances of their data are clear.
5. **Create compact interactive visualizations**: Interactive visualizations promote effective communication. `dms-viz` creates compact plots that can be incorporated into HTML presentation slides (e.g., https://slides.com/).
6. **Share findings with ease**: Users of `dms-viz` can generate shareable URL links for a customized visualization view. They can also save and share the `JSON` specification files created by the command line interface, ensuring that data can be accessed easily.
7. **Preserve data privacy**: `dms-viz` allows users to analyze proprietary and sensitive data by supporting local upload. This means researchers can view and analyze their confidential structures and datasets without the requirement to store them in a public repository.

Our group previously created a tool called `dms-view` (Hilton et al. 2020) that has some of the functionalities listed above. However, we designed `dms-viz` to be more customizable and comprehensive to handle a wider diversity of experimental designs and questions.

## Design and Usage

Using `dms-viz` involves three components. First, using the command line tool `configure-dms-viz`, available as a Python package on PyPI (https://pypi.org/project/configure-dms-viz/), the user formats their data into a `JSON` specification file. Then, the user uploads this specification file to `dms-viz.github.io`, a web-based interface written in Javascript, `D3.js`, and `NGL.js` (Rose et al. 2018). Finally, the specification file can either be shared directly or hosted remotely to generate a shareable URL link (Figure 2).

**Figure 1.**
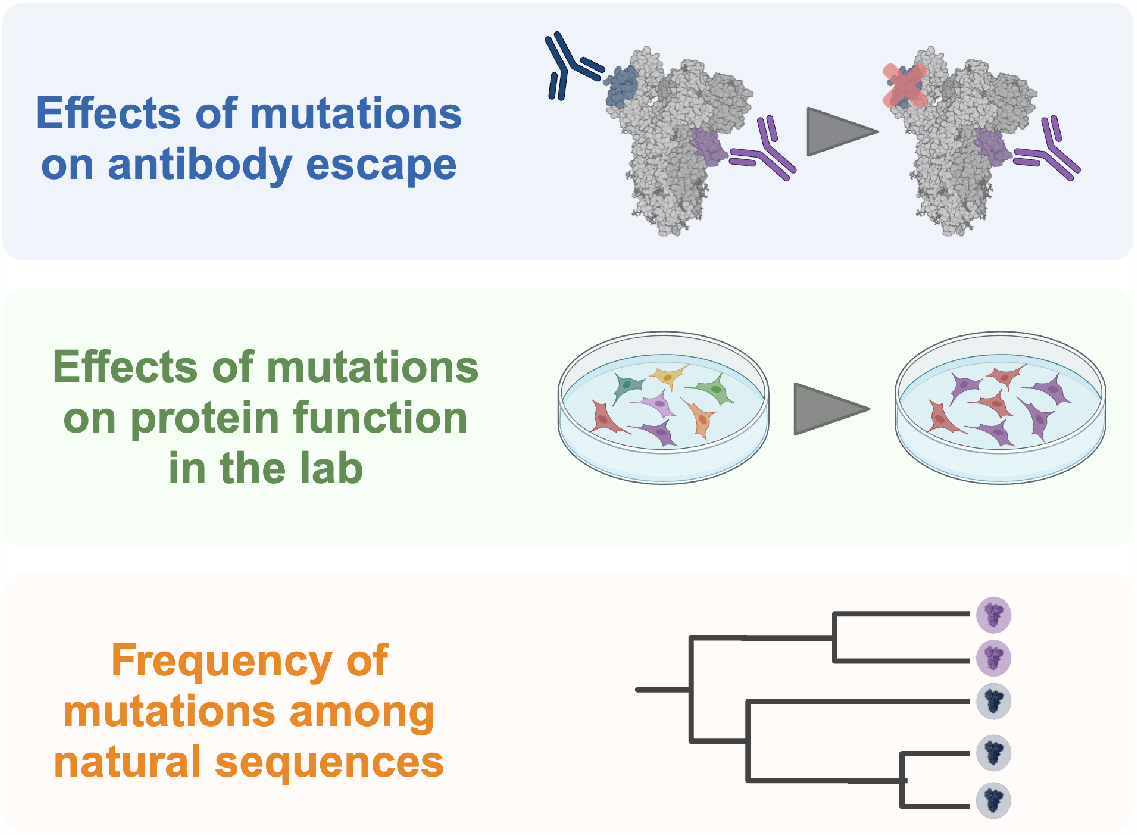
Large mutation-associated datasets are used in a variety of experimental contexts. They can be used to map antibody footprints on viral glycoproteins, assess the impact of mutations on protein function in a laboratory setting, and identify patterns of selection from natural mutation frequencies.

**Figure 2.**
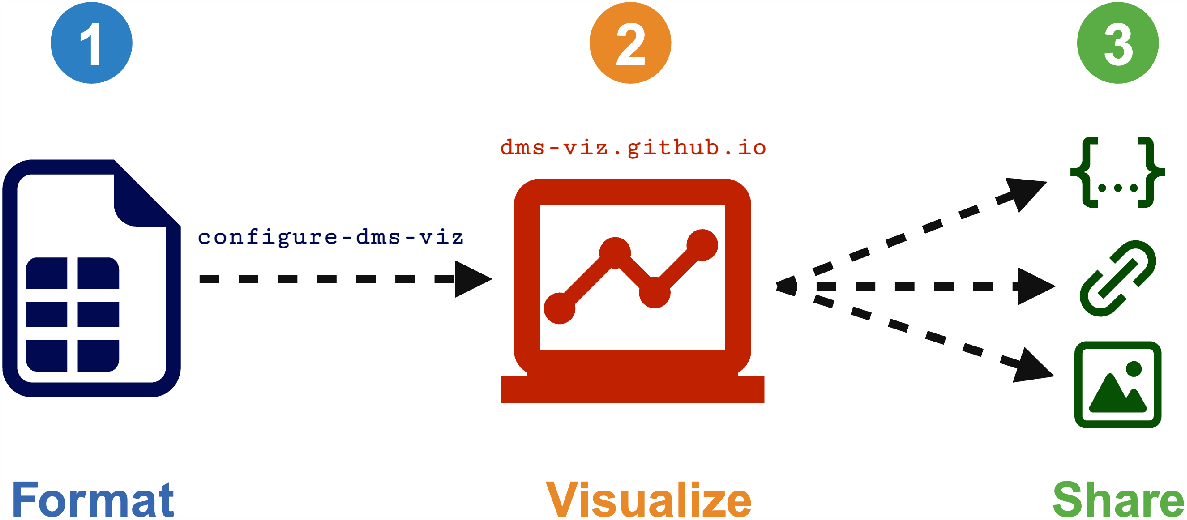
Using `dms-viz` involves three components. (1) The user formats their data using the command line tool `configure-dms-viz`. (2) The user takes the resulting `JSON` specification file and uploads it to `dms-viz.github.io`. (3) The user can choose to either share the `JSON` file, host the `JSON` file and generate a shareable URL link, or export static images.

Upon uploading the specification file to `dms-viz`, users will see a visualization composed of four components, as illustrated in Figure 3.

**Figure 3.**
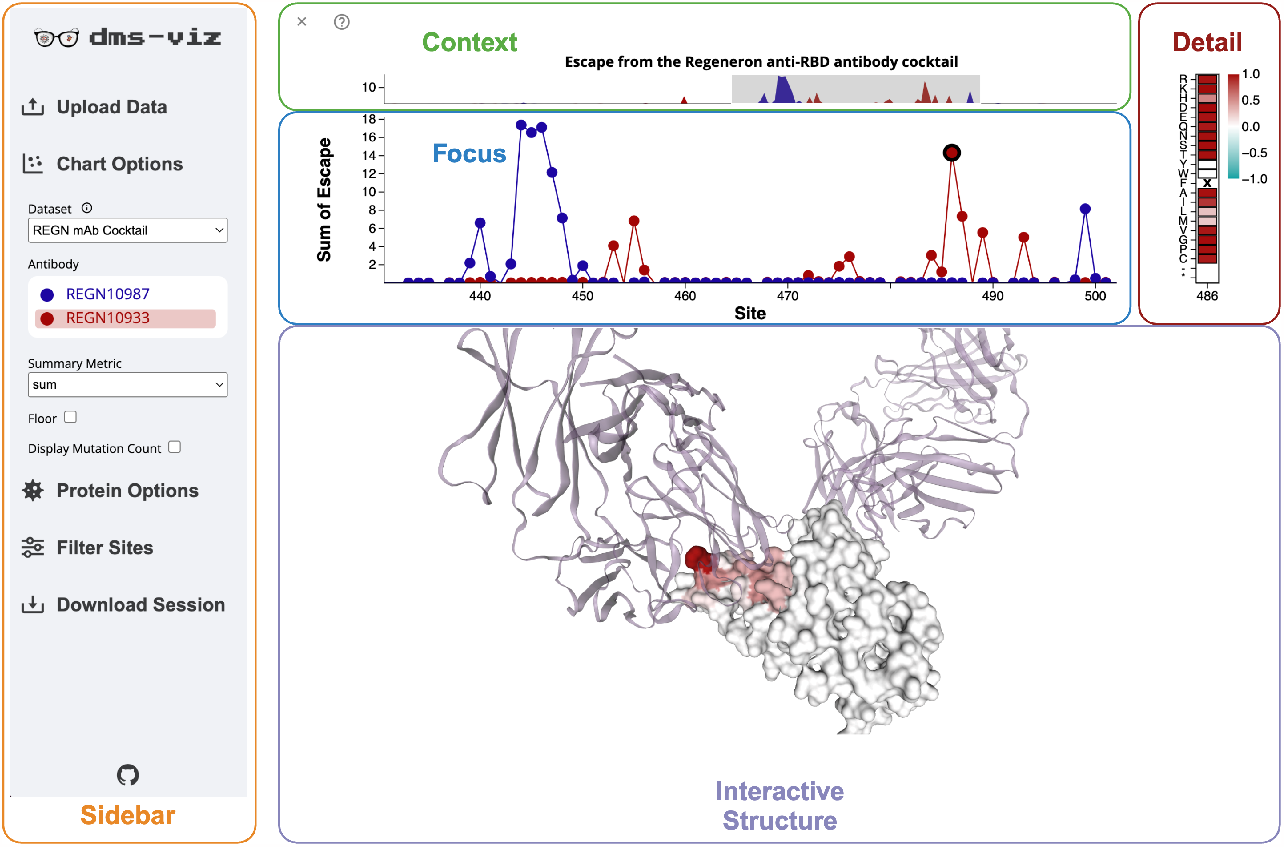
`dms-viz` provides a compact interface for exploring mutation-associated data. The visual component of `dms-viz` contains a line/point plot that shows a summary of the mutation-metric at all sites, in this case, mutation-escape from the constituents of a therapeutic antibody cocktail measured by DMS of the SARS-CoV-2 receptor binding domain (RBD) (Starr et al. 2021). The user can zoom into specific regions of interest while maintaining context of the whole dataset using the context plot. Additionally, users can click on points in the focus plot to get details on every mutation for each site in the detail plot. Finally, sites that are selected on the focus plot by dragging are shown on the interactive protein structure colored by the summary statistic. In this example, the structure shown is the SARS-CoV-2 RBD bound to both antibodies in the therapeutic cocktail (PDB: 6XDG). A collapsible sidebar is used to configure the visualization and select the condition on the interactive protein structure. By collapsing out of view, the sidebar makes the visualization an optimal size for integrating into online platforms like websites and HTML presentation slides.

1. ***Context* plot**: Located at the top of the visualization, this component allows users to zoom into specific sites on the *Focus* plot while maintaining an overview of the entire dataset.
2. ***Focus* plot**: This plot shows a summarized view of the user’s data. Every measured protein site is represented as a point providing a summary statistic of the effects of mutations at that site, and adjacent sites are connected with lines.
3. ***Detail* heatmap**: If the user is interested in the measurements for every mutation at a site, they can click on that site in the *Focus* plot. This will populate a heatmap with each individual mutation measurement at that site.
4. ***Interactive* structure**: When the user wants structural context for a given set of sites, they can drag a brush over the corresponding points in the *Focus* plot. This action will highlight those sites on an interactive 3D protein model.

To ensure the visualization remains compact, all configuration options are tucked away in a collapsible sidebar. See the documentation at https://dms-viz.github.io/dms-viz-docs/ for more information about how to use `dms-viz` along with detailed tutorials and examples.

## Examples

### 1. Mapping the neutralization profile of antibodies and sera against HIV envelope

Radford et al. (2023) mapped mutations to HIV envelope (Env) that affect neutralization by polyclonal human serum using a pseudotyping-based deep mutational scanning platform (Radford et al. 2023). One aim of their study was to examine how the sites of escape mutations related to HIV Env’s structure. `dms-viz` excels in generating these visualizations, especially for intra-experimental comparisons. Using `dms-viz`, it is possible to show multiple antibody footprints on a single summary plot.

See how `dms-viz` can be used to interactively visualize datasets with multiple conditions here.

### 2. Using mutation-fitness data to augment structure-guided drug design

Bloom and Neher developed a method to estimate the fitness effects of mutations to all SARS-CoV-2 proteins by analyzing millions of human SARS-CoV-2 sequences (Bloom and Neher 2023). These mutation-fitness estimates are useful for purposes such as attempting to design antiviral drugs that target functionally constrained sites where resistance is unlikely to emerge.

By merging Bloom and Neher’s data with structural views of a viral target like the SARS-CoV-2 main protease (Mpro) in complex with a bound ligand such as MAT-POS-e194df51-1 from the COVID Moon-shot project (Boby et al. 2023), `dms-viz` offers an intuitive way to visualize whether a ligand is targeting a mutationally tolerant binding pocket. Computational chemists can incorporate this information into the design process by screening for compounds that target sites where mutations have negative effects on viral fitness.

See how `dms-viz` can be used to enhance structure-guided drug design here.

### 3. Exploring the evolutionary potential of the influenza A polymerase PB1 subunit

The influenza RNA-dependent RNA polymerase (RdRp) is essential to viral replication, but little is known about the effects of mutations on RdRp function. To address this limitation, Li et al. (2023) measured the effects of thousands of mutations to the PB1 subunit of the RdRP on the replicative fitness of the lab-adapted influenza strain A/WSN/1933(H1N1) (Li et al. 2023).

`dms-viz` enables facile visualization of these data in the context of PB1’s structure, and can provide stable URL links for easy sharing and access.

See how `dms-viz` can provide this dataset as an interactive resource here.

## Conclusion

We designed `dms-viz` as a practical and user-friendly approach to visualizing mutation-associated data in the context of protein structures. Because `dms-viz` is capable of handling various data types and has options for both sharing and privacy, it should be applicable to visualization of a wide range of datasets.

## Code Availability

- `dms-viz` is available at https://dms-viz.github.io/
- The documentation and information about developing `dms-viz` is available at https://dms-viz.github.io/dms-viz-docs/
- The source code for `dms-viz.github.io` is available at https://github.com/dms-viz/dms-viz.github.io
- The source code for `configure-dms-viz` is available at https://github.com/dms-viz/configure_dms_viz

## Acknowledgements

This project was envisioned as the successor to the awesome tool `dms-view`. Thank you to Dr. Sarah Hilton and Dr. John Huddleston for laying this groundwork and for their incredibly helpful input on `dms-viz`. Thank you to members of the Bloom lab for providing data and guidance that was instrumental in developing `dms-viz`. Research reported here was supported in part by NIAID of the National Institutes of Health under award number U19AI171399. The content is solely the responsibility of the authors and does not necessarily represent the official views of the National Institutes of Health. The work of JDB was supported in part by the NIH/NIAID under grants R01AI141707 and contract 75N93021C00015. JDB is an Investigator of the Howard Hughes Medical Institute.

## References

Bloom, Jesse D., and Richard A. Neher. 2023. “Fitness Effects of Mutations to SARS-CoV-2 Proteins.” Virus Evolution 9 (2): vead055. 10.1093/ve/vead055.

Boby, Melissa L., Daren Fearon, Matteo Ferla, Mihajlo Filep, Lizbé Koekemoer, Matthew C. Robinson, The COVID Moonshot Consortium, et al. 2023. “Open Science Discovery of Potent Non-Covalent SARS-CoV-2 Main Protease Inhibitors.” bioRxiv. 10.1101/2020.10.29.339317.

Dadonaite, Bernadeta, Katharine H. D. Crawford, Caelan E. Radford, Ariana G. Farrell, Timothy C. Yu, William W. Hannon, Panpan Zhou, et al. 2023. “A Pseudovirus System Enables Deep Mutational Scanning of the Full SARS-CoV-2 Spike.” Cell 186 (6): 1263–1278.e20. 10.1016/j.cell.2023.02.001.

Hilton, Sarah K., John Huddleston, Allison Black, Khrystyna North, Adam S. Dingens, Trevor Bedford, and Jesse D. Bloom. 2020. “Dms-View: Interactive Visualization Tool for Deep Mutational Scanning Data.” Journal of Open Source Software 5 (52): 2353. 10.21105/joss.02353.

Li, Yuan, Sarah Arcos, Kimberly R. Sabsay, Aartjan J. W. te Velthuis, and Adam S. Lauring. 2023. “Deep Mutational Scanning Reveals the Functional Constraints and Evolutionary Potential of the Influenza A Virus PB1 Protein.” bioRxiv. 10.1101/2023.08.27.554986.

Radford, Caelan E., Philipp Schommers, Lutz Gieselmann, Katharine H. D. Crawford, Bernadeta Dadonaite, Timothy C. Yu, Adam S. Dingens, Julie Overbaugh, Florian Klein, and Jesse D. Bloom. 2023. “Mapping the Neutralizing Specificity of Human Anti-HIV Serum by Deep Mutational Scanning.” Cell Host & Microbe 31 (7): 1200–1215.e9. 10.1016/j.chom.2023.05.025.

Rose, Alexander S., Anthony R. Bradley, Yana Valasatava, Jose M. Duarte, Andreas Prlic, and Peter W. Rose. 2018. “NGL Viewer: Web-Based Molecular Graphics for Large Complexes.” Bioinformatics (Oxford, England) 34 (21): 3755–58. 10.1093/bioinformatics/bty419.

Starr, Tyler N., Allison J. Greaney, Amin Addetia, William W. Hannon, Manish C. Choudhary, Adam S. Dingens, Jonathan Z. Li, and Jesse D. Bloom. 2021. “Prospective Mapping of Viral Mutations That Escape Antibodies Used to Treat COVID-19.” Science (New York, N.Y.) 371 (6531): 850–54. 10.1126/science.abf9302.

Starr, Tyler N., Allison J. Greaney, Sarah K. Hilton, Daniel Ellis, Katharine H. D. Crawford, Adam S. Dingens, Mary Jane Navarro, et al. 2020. “Deep Mutational Scanning of SARS-CoV-2 Receptor Binding Domain Reveals Constraints on Folding and ACE2 Binding.” Cell 182 (5): 1295–1310.e20. 10.1016/j.cell.2020.08.012.

Yu, Timothy C., Zorian T. Thornton, William W. Hannon, William S. DeWitt, Caelan E. Radford, Frederick A. Matsen, and Jesse D. Bloom. 2022. “A Biophysical Model of Viral Escape from Polyclonal Antibodies.” Virus Evolution 8 (2): veac110. 10.1093/ve/veac110.

